# Electrostatic Fields Have Strong Repellency Effects Against *Culex pipiens*

**DOI:** 10.1101/2020.08.04.236828

**Authors:** Jonathan Tam

## Abstract

Currently, the most effective mosquito repellents employ the use of the chemical N, N-Diethyl-meta-toluamide (DEET), often considered the gold standard of repellents. However, as DEET continues to be widely used, it is beginning to lose efficacy as mosquitoes gradually gain resistance to it. Additionally, DEET only protects the skin that it is applied on, and therefore, it is difficult to achieve 100% repellency for the entire body. As urbanization and climate change continue to increase the threat of widespread mosquito-borne illnesses, new forms of repellents must be developed to combat the possibility of an epidemic. Previous studies indicate that a multitude of insects, including spiders, flies, and mosquitoes, avoid electrostatic fields at sufficiently high voltages. In this study, I evaluate if electrostatic fields can be used as a potential personal repellent against mosquitoes and an alternative to chemical sprays. I found that by charging the surface of a person’s body with a high voltage (9800V) but minuscule current, I created a safe and effective method that repels mosquitoes from landing on the skin. This method has similar efficacy to DEET, with near 100% repellency after 20 minutes of charging. It also has the advantage of not needing to be reapplied and provides full-body protection, allowing greater protection to the user than DEET. These results introduce a promising method of repelling mosquitoes, that used together with present-day chemical repellents, can help provide greater protection to malaria-endemic countries.

## Introduction

Mosquitoes are known to carry many deadly diseases, such as dengue, West Nile Virus, and chikungunya [1]. Malaria, one of the deadliest mosquito-borne diseases, caused 228 million cases in 2018, with the majority of these cases in the African and South-East Asia regions [2,3]. Access to current technology in insect repellents, vaccines, and drugs are severely limited in these regions, leading to a greater number of mosquito related deaths than other parts of the world [3,4]. In the Americas, dengue has also been a deadly disease spread by the *Aedes* family and resulted in 96 million cases annually, with cases continuing to rise in recent years [1,5,6]. Future predictions indicate that climate change and urbanization will increase the spread of mosquitoes at a quicker rate, increasing the risk of epidemics [7–9]. As a result, it is important that more effective methods continue to be developed.

There are two major types of repellents that are mainly used: topical and spatial [10]. Topical repellents, chemicals that are applied directly to the skin or clothing, include sprays that use deterrents such as DEET, picaridin, or lemon eucalyptus [11]. While these chemicals are generally seen as effective repellents, they require reapplication at regular time intervals to remain effective [12,13], making them too expensive and impractical for many places at high risk for mosquito-borne diseases [14,15]. Mosquitoes are also beginning to become resistant to DEET, which could possibly render the chemical ineffective in the future, increasing the risk of epidemics and consequently increasing the needs for other methods to be developed [16–19]. Examples of spatial repellents include citronella candles, which have had mixed result in effectiveness, but in general are not nearly as effective as topical repellents nor seem to provide 100% protection for any amount of time. [20–22]. There have also been several attempts to use electronic methods to repel mosquitoes using a high-pitched noise, also known as ultrasonic repellency, but none have been proven to have any legitimate effect [23].

Only been a few studies experimenting with whether electrostatic fields have any ability to repel insects [24–26]. These have mostly been focusing on its use in agriculture to protect crops from pests, leaving its potential application as a personal mosquito repellent unstudied, to my knowledge [27–29]. This phenomenon has also been found in *Drosophila*, as electric fields have also been found to induce avoidance behavior in them by affecting their wings, and the stronger the electric field, the greater the avoidance behavior observed [30]. Because electrostatic fields appear to have a prominent effect on *Drosophila’s* wings, the question of whether electrostatic fields may cause the same avoidance behavior in mosquitoes comes to mind. I aim to further explore the effects of electrostatic fields on insects, particularly mosquitoes, and evaluate its ability to be used by as a possible alternative to current repellents.

## Results and Discussion

### Electrostatic Fields Induce Avoidance Behavior in Mosquitoes

To understand how electrostatic fields affected mosquito blood-feeding behavior, a modified arm-in-cage test was conducted, where an entire exposed forearm and hand was offered to the mosquitoes (Figure 1A) [31]. While previous avoidance assays used a charged net to deter insects from passing through, I chose the arm-in-cage test to better compare electrostatic fields to current repellents [31, 32]. The attractant’s (test subject) skin was charged with voltages ranging from 1400V to 9800V, and the number of landings were observed for each voltage. This lower voltage range was based on previous assays with how other insects behaved (previous studies showed around 2kV charged nets were effective against the majority of tested insects) and the highest voltage to be tested was based on the limits of the electrostatic generator [25,27,33,34].

**Figure 1:**
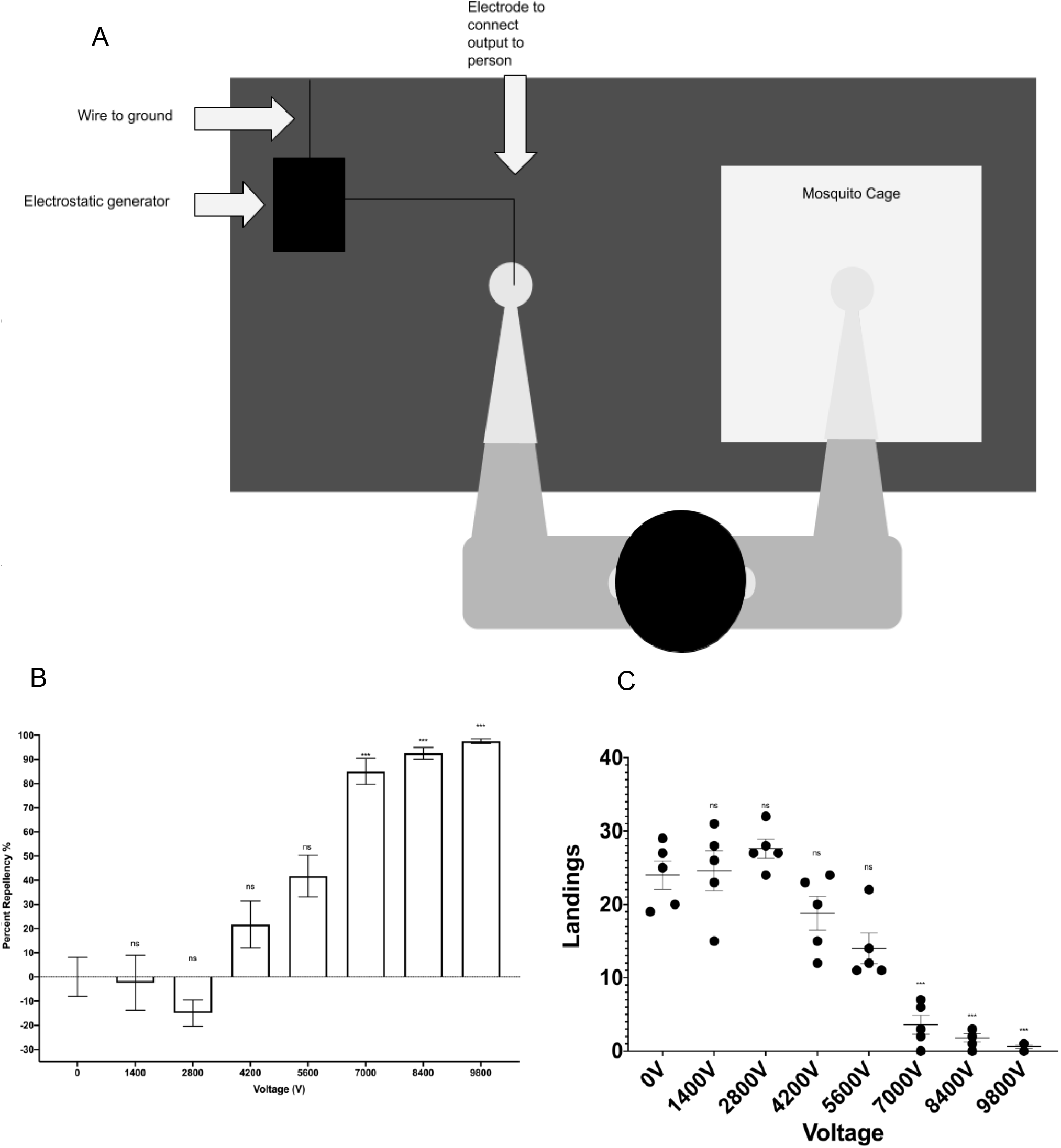
Electrostatic Fields Induce Avoidance Behavior in Mosquitoes. (A) Schematic of arm-in-cage test with electrostatic generator. The voltage can be changed from the electrostatic generator device, which charges the surface of the skin. (B) Repellency Rate for Voltage 0V-9800V (C) Number of Landings for Voltages 0V to 9800V Horizontal lines in (B-C) represent mean and ± SEM. “ns” indicate statistically insignificance, while *** indicates statistical significance, p < 0.0001 (one-way ANOVA with post hoc Tukey’s HSD test were used. Bars represent average repellency rate in (B). Dots represent individual trials in (C). N=5 experiments, 45-50 mosquitoes/test.

From 0V to 2800V, the repellency rate remained at 0%, and surprisingly, these voltages had practically no effect on the mosquito landing behaviors. In fact, more mosquitoes landed on average than without any charge, indicating that mosquitoes were not affected by these voltages (Figure 1B). This lack of effectiveness, even at 2800V, is most likely because electrostatic fields decay according to an inverse square law, meaning a higher voltage must be used. Beginning at 4200V, the repellency rate began to increase, and at 7000V, repellency reached 85% ± 5.37% (Figure 1B). At this voltage, the number of landings dropped significantly, from approximately 19 landings to under 4 (Figure 1C). This was the most significant increase in repellency, as the repellency rate increased by 43.3% at this voltage. At a voltage of 9800V, a maximum of 1 mosquito landed in a test (Figure 1C), leading to an average repellency rate of 97.48% ± 1.02% (Figure 1B).

These tests provide strong evidence that electrostatic fields do induce significant avoidance behavior in mosquitoes, especially at 9800V, and demonstrates that this method can be developed into a possible alternative form of repellent. As voltage increases, it is clear that mosquitoes exhibit increasingly prominent avoidance behavior, and this becomes significant at 7000V, demonstrating the existence of a “threshold” electrostatic field strength in which they begin to avoid the charge, confirming the experiments by Matsuda et al. [33]. At 9800V, the repellency rate and number of landings are comparable to the efficacy of DEET [11,35].

Compared to current commercial products, such as DEET or picaridin, electrostatic fields have proven to be as effective, if not more effective, with a near 100% repellency rate [11,21,35]. This method presents an advantage that chemical repellents do not have, in that it does not require reapplication, only requiring to have a constant power source. On the other hand, DEET requires reapplication every 6 hours in order to remain usable, one of the largest disadvantages of the repellent [21,36]. Additionally, DEET only protects the skin that it is applied on and can only be applied on exposed skin (cannot be covered by clothing); therefore, it is difficult to achieve 100% repellency for the entire body and leaves parts of the body vulnerable [36]. However, since static electricity builds up over the entire surface of the skin [37], electrostatic fields may be able to provide full body protection unlike chemical repellents. This is proven by the ability of the charges to move across from one hand (where the electrostatic generator contacts the skin) to the arm that was being tested (Figure 1A).

Some possible disadvantages of using this method is that the electrostatic generator must be grounded while the user is isolated from the ground (with shoes), which could cause inconvenience. Discharge is also probable, if the user touches other grounded objects.

### Repellency Rate Increases as the Time the Attractant is Charged Increases

While it is known that static electricity builds up on the outside of skin [38], little else is known about how charges behave on human skin. During the conception/initial experiments of this study, I observed that the longer the time the attractant was charged for, the fewer mosquitoes landed, even at the same voltage. Consequently, I found that charges took a significant amount of time to spread across the rest of the body, meaning this could potentially affect the efficacy of the electrostatic repellent. To further understand how the time charged affected mosquito behavior, I performed a modified arm-in-cage test.

This assay revealed that after a significant amount of time being charged, approximately 20 minutes, the repellency rate of the electrostatic fields remained at a high percentage and kept stable (Figure 2B). At a charging time of 0 minutes, with 9800V, the repellency rate remained at a relatively low 75.83% ± 2.43%, with about 6 landings, and the repellency gradually increased as the time the attractant was charged increased, eventually reaching 99% ± 0.83% repellency at 20 minutes and 100% at 25 and 30 minutes (Figure 2A-2B). No landings occurred from 25-30 minutes in any of the trials (Figure 2A). This is significant, as it indicates that the longer the electrostatic generator is worn, the greater the repellency effects. This test was only used for 9800V because it has been established in previous tests to have the highest repellency effects on the mosquitoes.

**Figure 2:**
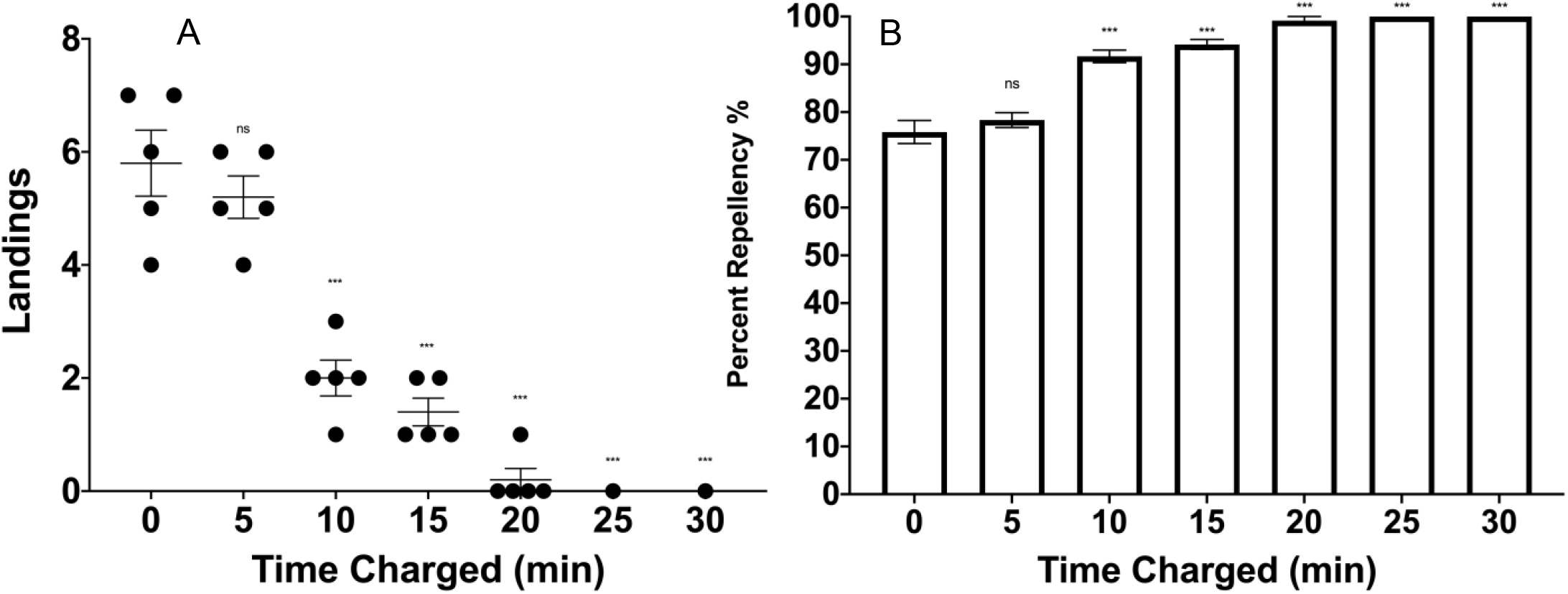
Repellency Rate Increases as the Time the Attractant is Charged Increases. (A) Number of Landings vs. Amount of time skin was charged. (B) Repellency Rate vs. Amount of time skin was charged. Horizontal lines in (A-B) represent mean and ± SEM. “ns” indicate statistical insignificance, while *** indicates statistical significance, p < 0.0001 (one-way ANOVA with post hoc Tukey’s HSD test were used. Dots represent individual trials in (A). Bars represent average repellency rate in (B). N=5 experiments, 45-50 mosquitoes/test.

Because the charges require time to migrate across the skin, it appears that method of repellency requires a “lead time” before being fully effective. If this method is used for repelling mosquitoes, it is important that it is wearable for long periods of time and effective at all times. Even though this requires a minimum 20 minutes before this method becomes fully effective, the long-term effectiveness means it can outlast any chemical repellent (as long as the electrostatic generator is powered). Unlike other repellents, this method increases in efficacy as time goes on, as demonstrated by the increase in repellency rates from 0-30 minutes (Figure 2A). For living conditions that require long-term mosquito protection, this method can potentially help solve the problem of chemical repellents being too expensive for low-income countries [14,15], as the cost over time of use would decrease.

In order for electrostatic fields to be used as a repellent, it necessarily exposes the person using it to the fields. While there have been concerns about the potential harmful effects of high electric fields on humans, these appear to be inconclusive or miniscule [37,39,40]. The only small risk of static electric fields are micro-shocks, which only cause a minor nuisance and have no real adverse effects [37,41]. However, throughout experimentation, even at the highest voltage of 9800V, there were no incidences of micro shocks, though further studies can be conducted to better understand the risks of micro shocks at relatively high voltages.

## Limitations and Future Work

While this study demonstrates that *Culex pipiens* are repelled by electrostatic fields, further studies must be done to test whether other families, such as the *Anopheles* and *Aedes*, will also be repelled in the same manner. Furthermore, future work could also involve researching the reasons behind this phenomenon, as an understanding of this could help develop a better solution to repelling mosquitoes. Developing a low-cost method of generating this high voltage will help those who are unable to afford chemical repellents, increasing the number of people who have access to personal insect protection. Future studies may also incorporate field testing to understand how electrostatic repellents fare in harsh weather conditions.

This study has solely focused on whether electrostatic fields can be used as a personal insect repellent, but has not done any work on whether this method can be used to with insecticides-treated nets, the main method used in malaria control [42]. As the number of usable insecticides is continuing to diminish as mosquitoes gain resistance [43], it is important that more strategies are employed to ensure the safety of these high-risk populations. Electrostatic fields could potentially be used in conjunction with insecticides-treated nets to further improve the protection of endemic nations. By charging these nets with an electrostatic fields, mosquitoes may be deterred greatly from approaching them, increasing the efficacy of the nets.

## Conclusions

With mosquitoes appearing to be an increasing threat as urbanization and climate change continues to change the world [6,44], and DEET, the most effective repellent as of now [16,21], appears to be losing efficacy, it is of great importance that new methods are used to counter this threat. While there have been attempts at solving this problem on a large scale, such as genetically modify mosquitoes to eventually render them unable to reproduce [45], this may also bring about unforeseeable consequences created by disrupting the global ecosystem [46], making it a questionable solution. For the time being, developing more effective repellents, such as the one described in this study, should still continue to be pursued.

Electrostatic fields have proven to provide substantial protection to the user, and outperforms many other mosquito repellents currently on the market. The work done in this study can serve as the starting point for further development of repellents that use electrostatic techniques to curb the effects of mosquito-borne illnesses on the world.

## Supporting information

Raw Data

## Materials and Methods

### Mosquito and Larvae Rearing

*Culex pipiens*, San Mateo strain, adult mosquitoes and larvae were obtained from the San Mateo County Abatement District and the Alameda County Abatement District. The adult mosquitoes were reared using procedures outlined by Kauffman et al. [47]. The ratio between male and female adult mosquitoes was maintained at 1:10, adult mosquitoes were fed 10% sucrose solution. Larvae were given powdered Wardley Fish Food Flakes, ad libitum, and pupae were moved to a small container in 30 x 30 x 30 cm BioQuip Cages. The mosquito and larvae environment were maintained at a temperature of 25-30°C and a relative humidity of 50%. The photoperiod maintained was 12:12 (L:D) h. No blood meal was provided, as *Culex pipiens* have the ability to reproduce without it [47]. The mosquitoes were allowed to mature for a minimum 5 days and starved for 12 hours before being tested upon.

### Electrostatic Generator

The electrostatic generator was constructed using a step-up transformer to increase the voltage to 700V and a voltage multiplier consisting of alternating diodes (1N4007) and capacitors (445-181678-ND) to increase the voltage to 9800V (Figure S1). Due to the incremental nature of the voltage multiplier, the output can be drawn from voltages 1400V-9800V in increments of 1400V (Figure S1), allowing for testing of various voltages. The circuit was contained within a plastic box (9.5 cm (L) x 3.5 cm (H) x 5.5 cm (W)), with the negative terminal grounded. An alligator clip was connected to the output and the opposite side was held by the attractant, charging the surface of the attractant’s skin [37]. The attractant wore shoes to ensure discharge did not occur. Note: Because of the design of the voltage multiplier, as voltage increases current decreases [48], thus allowing for a low enough power to be safe.

### Human Subject

The human subject used in this study was me (the author), and the procedures used were reviewed and approved by the Society for Science and the Public Institutional Review Board. I used the control experiments described in method details to determine that my body odor was attractive enough for mosquitoes.

## Method Details

### Testing of repellency effects of electrostatic fields

The arm used in testing was washed with unscented soap and dried. The electrostatic generator was turned on and held by the attractant for 20 minutes, building up static charge on the skin. The arm was then inserted into the cage of 45-50 mosquitoes (with an approximate 1:10 male to female ratio) for a total of 3 minutes per test, with the number of landings counted. The number of bites were considered irrelevant, as the goal was to investigate whether the mosquitoes avoided approaching and landing on the arm, and therefore were not counted.

Testing began a 0V (control) and continued in increments of 1400V until 9800V was reached. The experiments were completed every other night in early evening, as this was the most active time for *Culex pipiens* [49,50]. One voltage was tested every night, with 4 replicate tests performed at every voltage level, for a total of 5 tests. A control experiment was done before any testing to ensure the mosquitoes were active. For these tests, if 10 mosquitoes landed within a minute, the mosquitoes were considered active, and the rest of experimentation proceeded. Landings were counted manually and recorded at the end of each test. Only one attractant (test subject) was used throughout the testing.

### Finding the relation between time charged and avoidance behavior

Since static charges are known to build up on the surface of skin as time progresses [38], it was important to find the amount of time for the electrical charge to fully spread across the body. The procedure is the same as the previous section with the following modifications:

The arm is placed in as soon as the electrostatic generator is turned on, as opposed to being charged for 20 minutes. The arm is kept in the cage for 3 minutes and the landings counted, as previously done. It was then removed from the cage for 2 minutes, and then reinserted, summing up to a 5-minute time interval. During this time, the electrostatic generator was kept on to ensure that the attractant is always charged. This process was repeated for a total of 30 minutes. The attractant did not touch anything for the duration of the test to avoid the possibility of discharge.

## Quantification and Statistical Analysis

The efficacy of each particular voltage was evaluated using one-way analysis of variance, with the independent variable being the voltage and the dependent variable being the number of landings. Statistical significance was defined as p < 0.05. The repellency rate was found by subtracting the number of landings for a given voltage from the number of control landings and dividing by the average number of control landings, multiplied by 100 [51]. This provides a percent repellency that can be compared with the control and other voltages.

Statistical analysis was done using SPSS and GraphPad Prism 8. Details for how software was used, such as statistical tests and significance, can be found in figure legends. All raw data is available in Data S1.

## Acknowledgements

I would like to thank the San Mateo Mosquito Abatement District and the Alameda County Mosquito Abatement District, for generously providing me with the mosquitoes.

