## Supplemental Figure 1 for "Electrostatic Fields Have Strong Repellency Effects Against *Culex pipiens*"

### Supplemental Information

### Data:

**Data S1:** Raw Data can be found in the excel file titled “Supplemental Data (Data S1)”

### Figures:

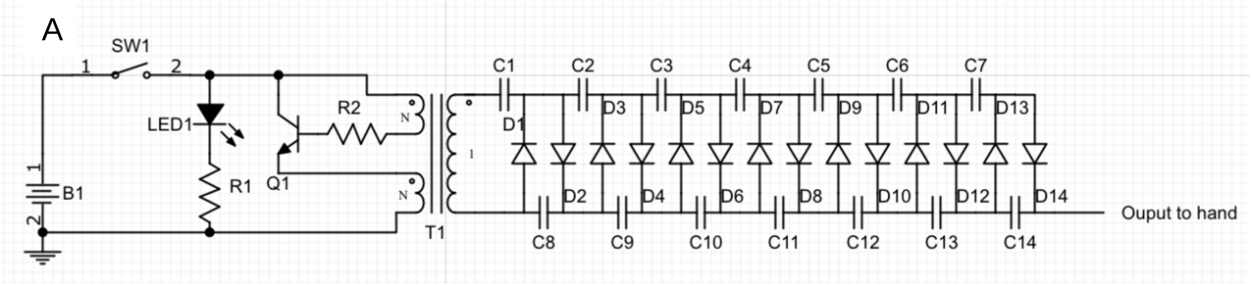


**Figure S1: Electrostatic Generator**

(A) This generator was created with step-up transformer in series with a voltage multiplier that allows the voltage to be stepped up to 9800V. By placing an alligator clip electrode in between two capacitors, a lower voltage can be generated. For example, between C8 and C9, a voltage of 1400V is outputted. This allows for the incremental increase in voltage.

Parts List:
B1: 9V Battery
LED1: LED Light
R1: 1 kΩ
R2: 2.2 k kΩ
SW1: Switch
Q1: D965
T1: EE19
C1-C14: 445-181678-ND
D1-D14: 1N4007
